# Structural characterization of hibernating ribosomes in four Gram-negative pathogenic bacteria

**DOI:** 10.1101/2025.10.07.680626

**Authors:** Vasanthakrishnan Radhakrishnan Balasubramaniam, Olivier Delalande, Sophie Chat, Yann Lefrançois Copy, Sylvie Georgeault Daguenet, Mohamed Sassi, Emmanuel Giudice, Reynald Gillet

## Abstract

Ribosomes are universal molecular machines that translate messenger RNA into proteins. Depending on what the cell’s needs for protein synthesis, ribosomes exist in various functional states, and when cells face stressful conditions, their ribosomes may enter into an inactive hibernation state. In bacteria, this ability to become dormant through ribosomal hibernation plays a crucial role in their survival, especially when the cells are targeted by ribosome-targeting antibiotics. The phenomenon occurs when ribosomes bind with hibernation factors. In this work, we examine how these special proteins interact with ribosomes from four different Gram-negative pathogens, and how this leads to translation inhibition, reporting the single-particle cryo-electron microscopy structures of hibernating ribosomes from *P. aeruginosa, E. hormaechei, K. quasipneumoniae,* and *A. baumannii*. We show that these ribosomal complexes contain either short HPF or YfiA tightly bound to the 30S A-site, P-site, and even occasionally to the E-site. Our results provide valuable insights into the ribosomal hibernation mechanism, paving the way to both a better understanding of bacterial dormancy and the possibility of developing antibiotics which will target hibernating ribosomes rather than just active ones.

## INTRODUCTION

Cells spend a substantial portion of their energy on ribosome driven protein synthesis, reflecting the important role of translation in cellular growth and adaptation. To survive, they need to respond quickly to changing environments, so this process is tightly controlled (Warner, 1999). When cells adapt to unfavorable conditions, protein synthesis is retooled for energy and resource conservation until more favorable conditions arise (Warner, 1999; Njenga *et al*, 2023; Yin *et al*, 2019; Serbanescu *et al*, 2020). A key strategy for achieving this essential transformation is the use of hibernating ribosomal complexes, molecular factors which then place the active ribosomes into a dormant state (Prossliner *et al*, 2018). Ribosome hibernation has been observed under a wide variety of stress conditions, including nutrient deprivation (stationary phase growth) (Aiso *et al*, 2005; Maki *et al*, 2000), host colonization (Kline *et al*, 2015), adaptation to darkness (Contreras *et al*, 2018; Tan *et al*, 1994), temperature shock (Ranava *et al*, 2022; Helena-Bueno *et al*, 2024), amino-acid starvation (Izutsu *et al*, 2001), and biofilm formation (Williamson *et al*, 2012). This dormancy has been seen in all three domains of life, emphasizing its evolutionary significance (Trösch & Willmund, 2019), and its presence in widely diverse biological systems indicates that it is critical for cellular stress-response and survival.

Ribosome hibernation involves hibernation factors, and these belong to specialized families of proteins which are involved in bacterial survival under various stress conditions. In bacteria, researchers have so far characterized the structures of three of these families of ribosomal complex-forming factors (Figure 1 & Supp. Figure 1): ribosome modulation factor (RMF; Prossliner *et al*, 2018; Polikanov *et al*, 2012; Yoshida *et al*, 2004; Wada *et al*, 1995); the HPF/RaiA family containing the hibernation-promoting factor and/or ribosome-associated inhibitor A (Polikanov *et al*, 2012; Prossliner *et al*, 2018; Ueta *et al*, 2008; Sato *et al*, 2009; Ueta *et al*, 2013; Maki *et al*, 2000); and the more recently described Balon family (Helena-Bueno *et al*, 2024). When acting sequentially in *gamma-*proteobacteria, RMF and the short form of HPF (SHPF, previously known as YhbH) facilitate the dimerization of 70S ribosomes and the stabilization of 100S ribosomes (Figure 1A and 1B; (Ueta *et al*, 2005; Yoshida & Wada, 2014). In *Escherichia coli,* for example, RMF promotes the formation of a 90S complex, and then 90S ribosome dimer matures with the addition of SHPF to form 100S hibernating dimers (Ueta *et al*, 2005; Maki *et al*, 2000). In addition, *Gamma-*proteobacteria also possess a second hibernation promoting factor, protein Y (known as RaiA or YfiA). YfiA does not allow for 100S formation, and instead it binds directly to the 70S ribosomes to stop the translation activity of the monomeric form (Figure 1D; (Agafonov *et al*, 1999; Vila-Sanjurjo *et al*, 2004)).

**Figure 1.**
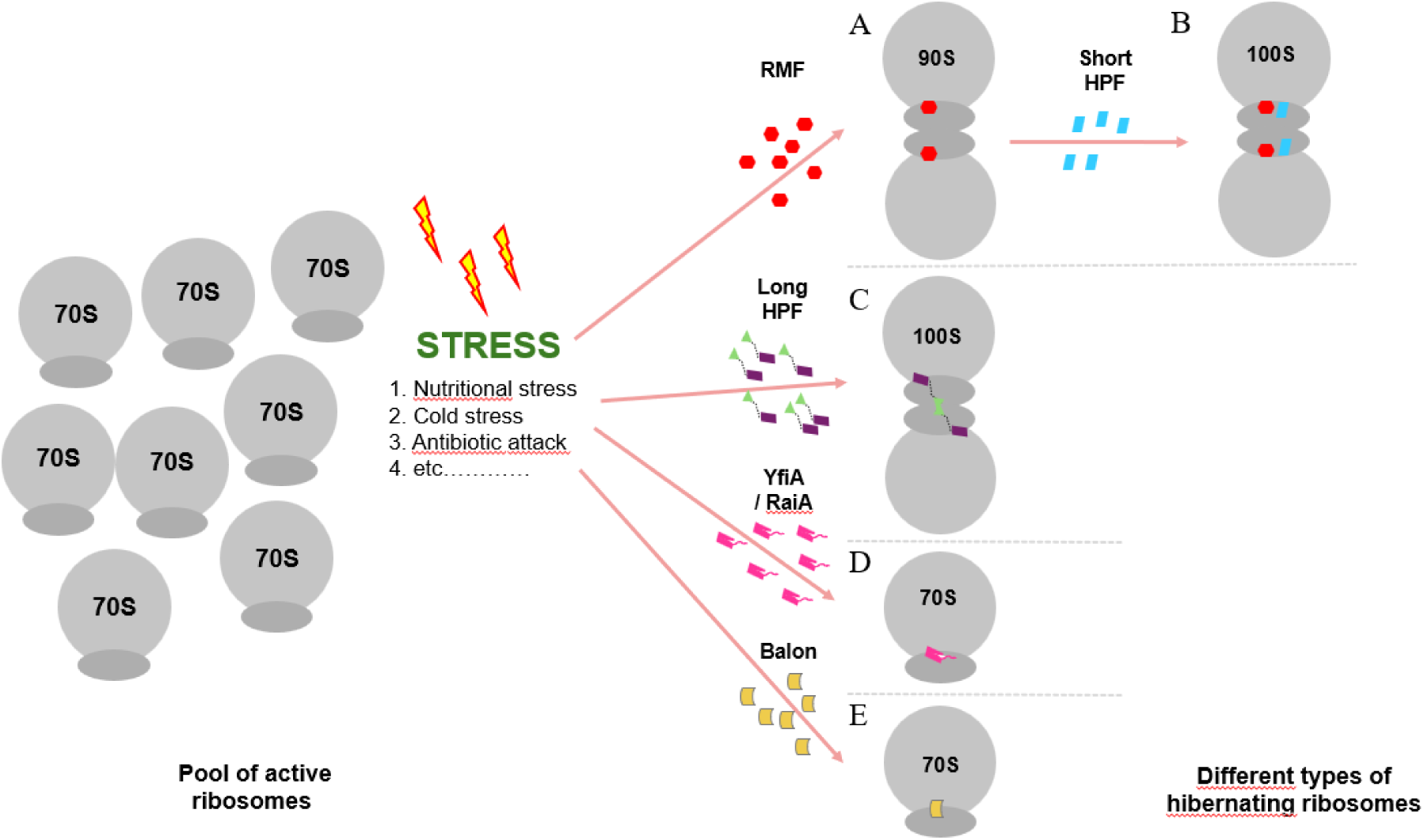
Schematic representation of ribosome hibernation mechanisms as a result of different hibernation factors in bacteria. Shown are hibernating ribosomes mediated by the proteins (A-B) RMF (red) and short HPF (blue); (C) long HPF (purple and green); (D) YfiA (also known as RaiA, magenta); and (E) Balon (yellow).

In other types of Proteobacteria, there is no RMF homolog, and these bacteria instead have a longer form of HPF (LHPF) which promotes 100S dimers (Figure 1C). In fact, recent phylogenetic analyses have revealed the presence of LHPF in almost all bacterial species except *gamma-*proteobacteria and *beta-*proteobacteria, with RMF and the shorter SHPF found only in a subset of *gamma-*proteobacteria, which includes *E. coli* (Gohara & Yap, 2018; Prossliner *et al*, 2021; Ueta *et al*, 2010, 2024; Akiyama *et al*, 2017; Prossliner *et al*, 2018). Sequence comparison studies of LHPF have revealed that its N-terminal domain is identical to that of the shorter SHPF, but that its C-terminal domain exhibits only a weak homology with RMF (Flygaard *et al*, 2018). The two LHPF domains are separated by a flexible region that often lacks a well-defined structure. In several bacterial species, including *Lactococcus lactis* (Franken *et al*, 2017), *Staphylococcus aureus* (Ueta *et al*, 2010), *Thermus thermophilus* (Flygaard *et al*, 2018), and *Bacillus subtilis* (Beckert *et al*, 2017; Tagami *et al*, 2012), LHPF is sufficient for the formation of 100S dimers without RMF. Such findings indicate that the mechanism of ribosome hibernation may vary across bacterial species, reflecting evolutionary divergence in the regulation of translational activity.

Over the past decade, developments in structural biology have extended our knowledge of ribosomal structures, providing a solid basis for understanding how ribosomes vary in composition across species and cellular compartments. Although we had structural details of a common ribosomal core, we lacked structural evidence of hibernating ribosomes in many important pathogenic bacteria. In this study, we provide such evidence for the four Gram-negative members of the ESKAPE family, a group of nosocomial pathogens presenting alarming levels of antibiotic resistance: *Klebsiella quasipneumoniae (K. pneumoniae complex); Acinetobacter baumannii; Pseudomonas aeruginosa P14 (“P. aeruginosa”* in the text); and *Enterobacter hormaechei subsp. Xiangfangensis* (*Enterobacter spp.,* hereafter *“E. hormaechei”*) (Tacconelli *et al*, 2018; Denissen *et al*, 2022; Mulani *et al*, 2019). The resulting detailed structural information about ribosome hibernation will help us to better understand cellular stress responses and protein synthesis regulation. In turn, this should aid in developing medical and biotechnological tools such as strategies for combatting bacterial persistence, as well as the improvement of antibiotic efficacy against these highly pathogenic bacteria.

## RESULTS

### Structures of hibernating ribosomes from *P. aeruginosa, E. hormaechei, K. quasipneumoniae,* and *A. baumannii*

We cultured ESKAPE bacteria to the early log phase, collecting the bacterial pellets after a final round of centrifugation at 4 °C, a temperature condition known to favor ribosome hibernation as part of the cell’s cold shock response (Agafonov *et al*, 2001; Helena-Bueno *et al*, 2024; Tissières *et al*, 1959). Remarkably, we found that significant populations of 70S ribosomes were in fact hibernating: 8% in *P. aeruginosa* and *E*. *hormaechei;* 14% in *A. baumannii;* and 26% in *K. quasipneumoniae* (Supp. Table 1). To further investigate the features of these hibernating ribosomes, we solved their structures, resulting in four cryo-EM maps at final resolutions between 2.5 Å and 2.8 Å (Figure 2). The next step was to clarify which hibernating proteins best fit into these density maps, so we began with a phylogenetic analysis of the *yfiA* and *hpf* genes (Figure 3). The resulting phylogenetic trees exhibit similar clustering patterns, highlighting the shared evolutionary relationship of the ESKAPE pathogens. In both trees, *E. hormaechei, K. quasipneumoniae,* and *E. coli* form a well-supported separate clade, suggesting a conserved evolutionary trajectory for these species. However, notable differences emerge in the placement of *P. aeruginosa* and *A. baumannii.* While *A. baumannii* strains (ATCC BAA-1605, ATCC 19606) form a separate lineage in both trees, the *P. aeruginosa* genome encodes only *hpf*, and it diverges significantly from the other species. This suggests that the *P. aeruginosa hpf* gene may have undergone greater evolutionary diversification, with functional adaptations perhaps triggered by unique circumstances. Overall, the comparative analysis reveals that while *yfiA* and *hpf* share conserved phylogenetic patterns across closely related species, their divergence increases among more distantly related ones, reflecting varying evolutionary pressures. While the other three bacteria contains both *yfiA* and *hpf* genes, *P. aeruginosa* harbors only the *hpf* gene. In our ribosomal structures containing both genes, the resolution was high enough to clearly distinguish the sequence and positional differences of specific residues of the two resulting hibernation factors: SHPF (hereafter referred to as ‘HPF’) and YfiA. Our analysis confirms that *E. hormaechei* contains HPF (Supp. Figure 2 A), while *K. quasipneumoniae* and *A. baumannii* contain YfiA (Supp. figure 2 B-C).

**Figure 2.**
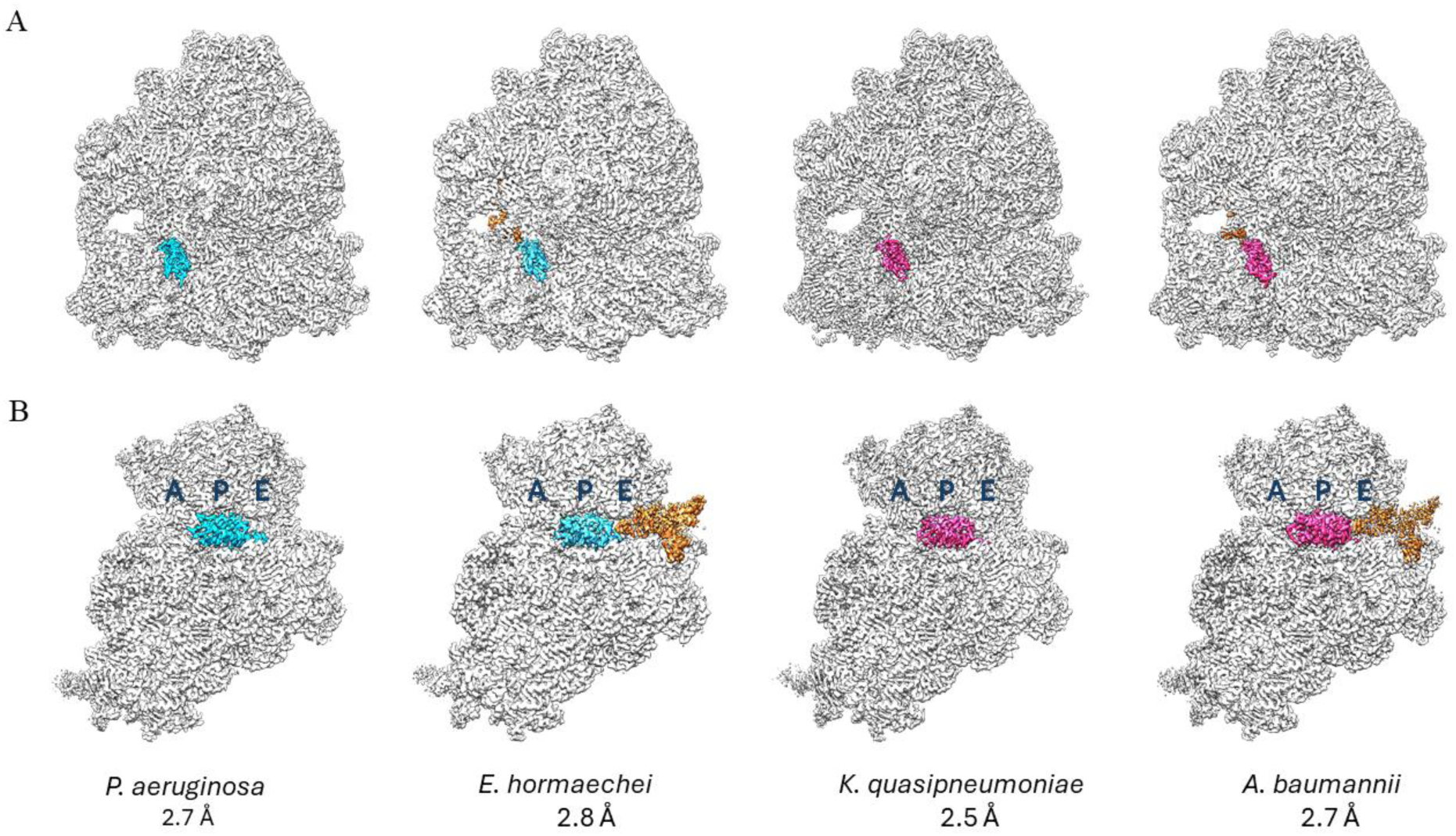
Structures of hibernating ribosomes in four Gram-negative ESKAPE pathogens. Top row (A), cryo-EM maps of 70S hibernating ribosomes. Bottom row (B), focus on the 30S ribosomal subunit showing the hibernation factors tightly bound on the mRNA channel between the head and body of the 30S, and overlapping the A- and P-sites (marked in B). HPF is cyan, YfiA is magenta, E-site tRNA is orange, and ribosomes are gray. Here we represent the most populated class of the maps (see text for detailed description).

**Figure 3.**
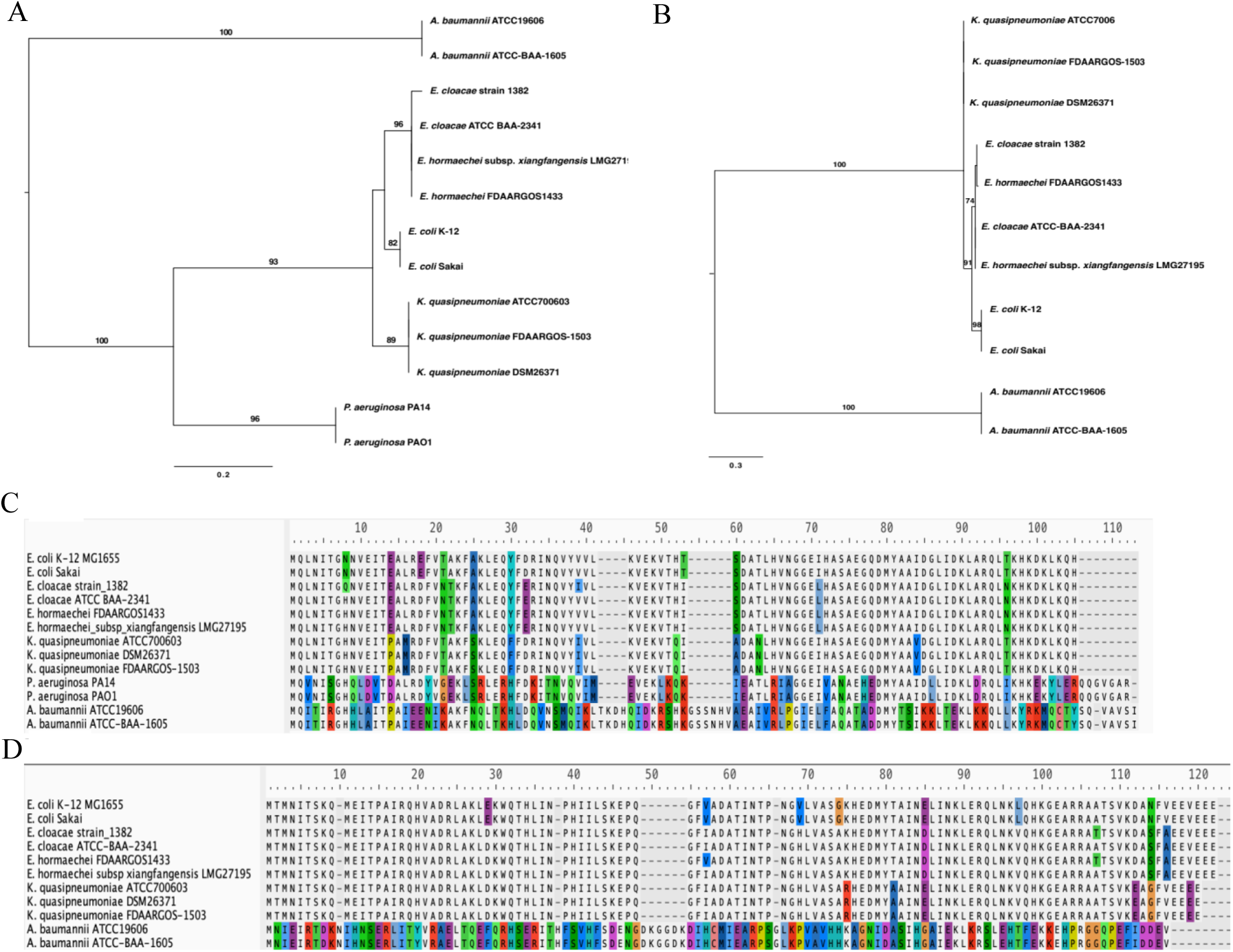
Phylogenetic analyses of ribosome hibernation-promoting factors in Gram-negative ESKAPE pathogens. Phylogenetic trees were constructed based on the protein alignments of (A) *hpf* and (B) *yfiA* genes using the maximum likelihood method. The bootstrap values (> 70%) were calculated for 1000 replicates and are shown at the bottom of each branch. Multiple sequence alignments for (C) *hpf* and (D) *yfiA* show sequence divergence across species. The conserved sites are highlighted, and amino acid substitutions, insertions, and deletions are indicated with standard symbols. The scale bar represents the relative genetic distance between taxa.

**Table 1.**
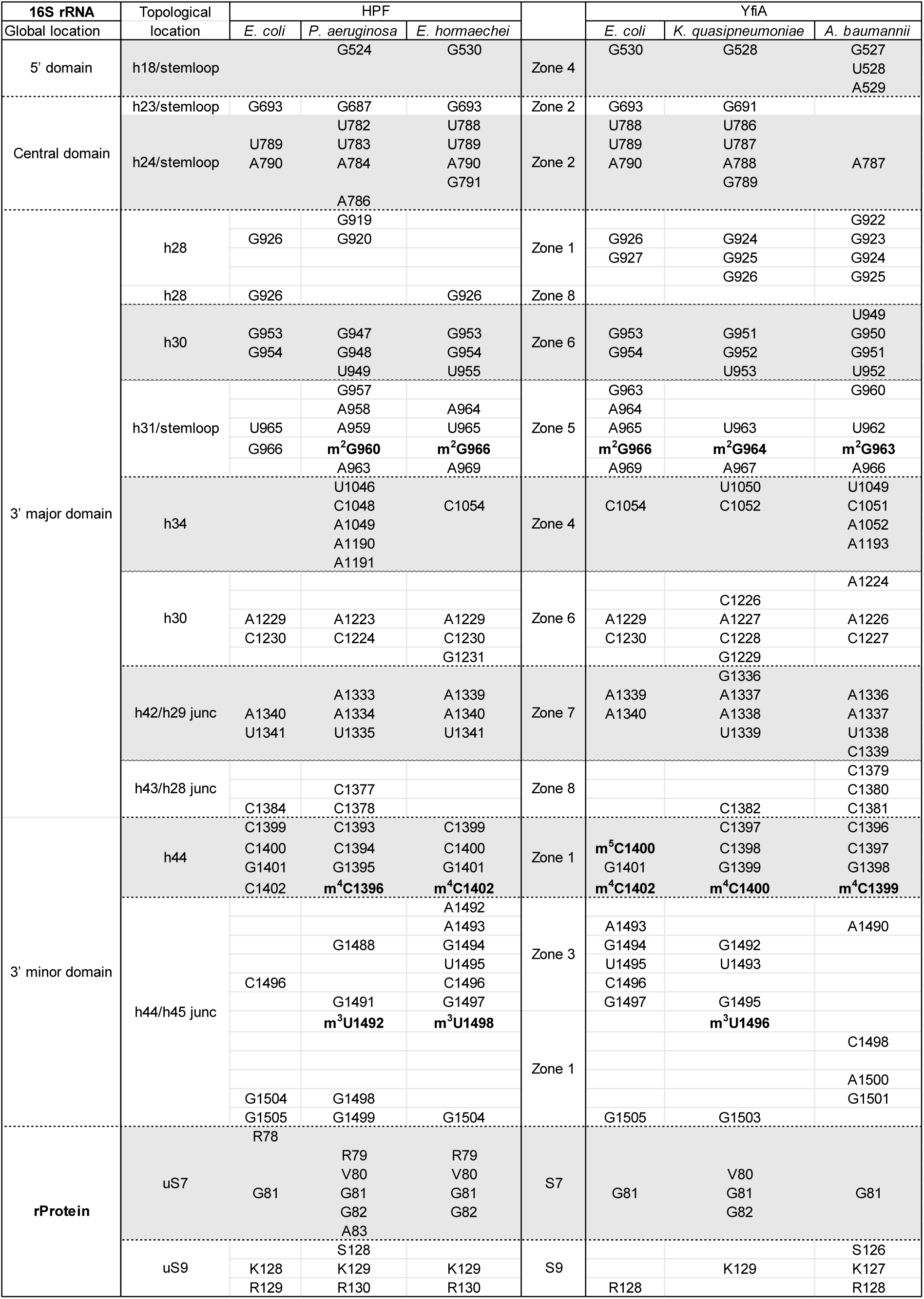
Summary of the ribosome nucleotides (16S rRNA) and amino acids (uS7 and uS9) in contact with hibernation factors HPF and YfiA. For the nucleotides, the global location within the 16S rRNA is listed in the first column, the detailed topological location is listed in the second column, and the corresponding representative zones (see Figures 4, 5, 7 and 8) in the sixth column. The numbering is specific to each pathogen studied here, with *E. coli* (PDB 6y69 for HPF, 8fc1 for YfiA) included as a reference. Modified nucleotides are in bold.

### HPF-ribosome interactions in *P. aeruginosa* and *E. hormaechei*

The structure of the HPF protein is composed of two α–helices and a four-stranded β-sheet (Supp. Figure 1). The HPF residues which interact with the ribosome are mainly located on the second α helix and on the β-strands. In the 70S-HPF complex structures we observed, the HPF protein binding is at the interface between the 30S and 50S subunits in both *P. aeruginosa* and *E. hormaechei*, yet it is mostly associated with the 30S at the A- and P-sites, and to a smaller degree, at the E-site (Figure 2B). In these two bacteria, HPF interacts with the 16S rRNA (mostly with the 3’ major and 3’ minor domains) and with the uS7 and uS9 ribosomal proteins (Table 1). The HPF interactions with the C-terminal end of uS9 are achieved through the conserved residues located at the end of β3 and at the beginning of β4 (see zone 7 in Figures 4, 5, and Supp. Figures 9 A-B and 12). As for the bS7 protein, it contacts the C-terminal end of HPF in both *P. aeruginosa* and *E. hormaechei* (see zone 8 in Figures 4, 5, and Supp. Figures 10 A-B and 12). The interactions between HPF and 16S rRNA are also very well conserved in these two bacteria, and mainly involve the helices h24 and h31 and their stem loops as well as helices h28, h30, h44, and the h42/h29 and h44/h45 junctions (Table 1). More specifically the HPF protein has contacts with several continuous tracks of nucleotides bearing post-transcriptional modifications, including m^2^G960, m^4^C1396, m^3^U1492 in *P. aeruginosa* (Figure 4 zones 1, 5, and Supp. Figure 12A), and m^2^G966, m^4^C1402, and m^3^U1498 in *E. hormaechei*. (Figure 5 zones 1, 5, and Supp. Figure 12B).

**Figure 4.**
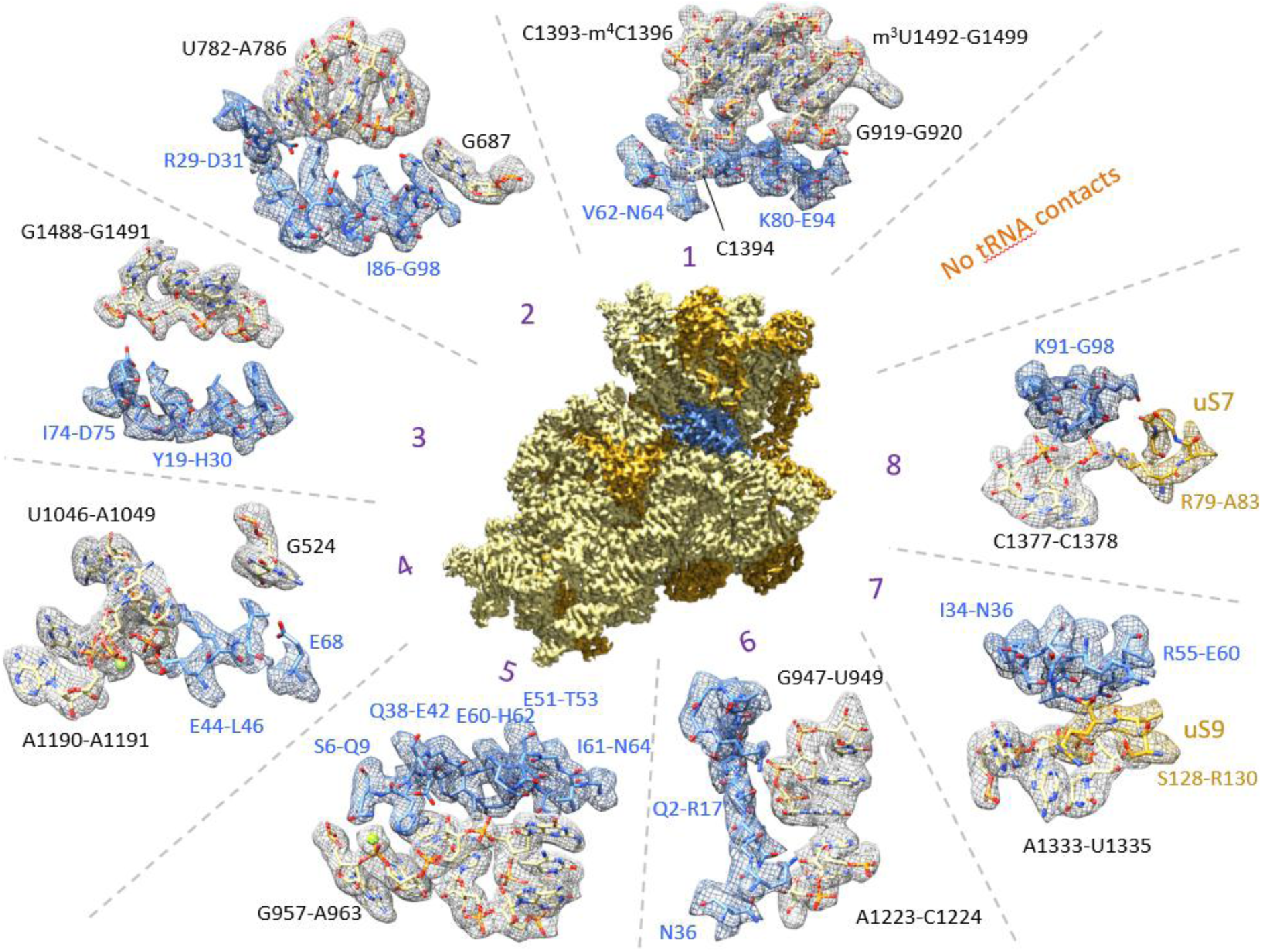
Interactions of the hibernation factor HPF with the *P. aeruginosa* ribosome. To aid in visualization, the interactions between the ribosomal small subunit 30S and HPF have been partitioned into geometric zones numbered 1-8. The nucleotide tracks of ribosomal 16S rRNA involved in binding to HPF are black, HPF-contacting amino acids are blue, ribosomal proteins uS7 and uS9 are gold, and tRNA nucleotides are orange.

**Figure 5.**
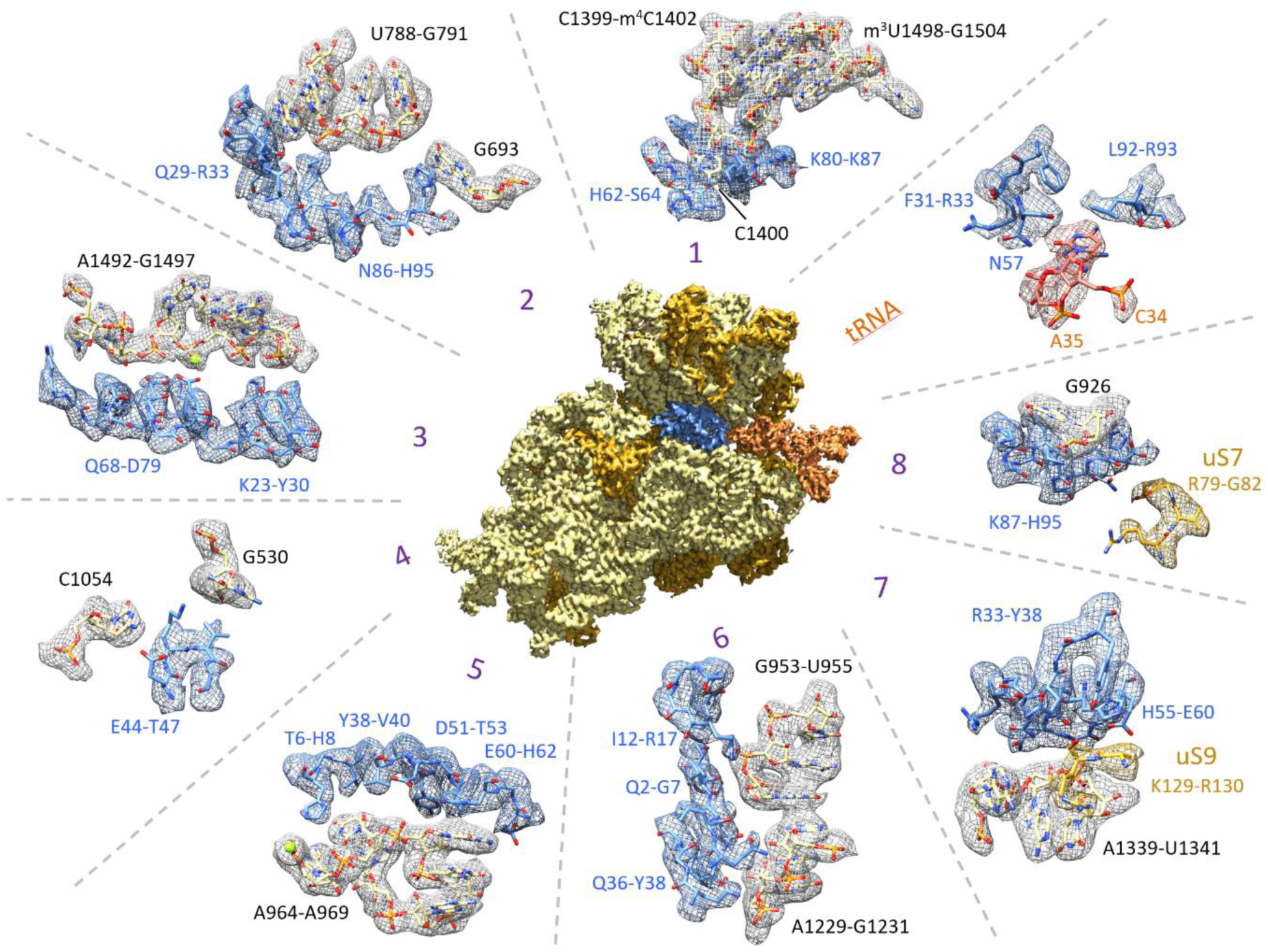
Interactions of the hibernation factor HPF with the *E. hormaechei* ribosome. To aid in visualization, geometric partitions are presented here by zones numbered 1 through 8 along with an addition tRNA zone (orange) to highlight the interaction between HPF and the E-site tRNA. The nucleotide tracks of ribosomal 16S rRNA involved in binding to HPF are black, the amino acids contacting HPF are blue, ribosomal proteins uS7 and uS9 are gold, and tRNA nucleotides are orange.

Even if the residues that interact with the ribosome are mainly located on secondary structural elements, we also identified some turns and/or loops which also carry interacting residues. In *P. aeruginosa* HPF, these include His8 and Gln9 (β1-α1 loop), Lys45 and Leu46 (β2-β3 loop), and Ala57 and Gly58 (β3-β4 loop) (Supp. Figure 12 A), while in *E. hormaechei* HPF these are found in His8 (β1-α1 loop), Glu32 and Arg33 (β1-β2 loop), Glu44 and Val46 (β2-β3 loop), and Asn57 (β3-β4 loop) (Supp. Figure 12 B). Similarly, the last residues of the HPF C-terminal extremity (such as His95 in *E. hormaechei*) also make strong contacts in both bacteria, even though they no longer belong to well folded α2 helix (Supp. Figure 12 A-B).

The HPF proteins from *P. aeruginosa* and *E. hormaechei* have nearly equal lengths (102aa and 95aa respectively, Figure 6) and occupy identical sites on the ribosome, so we detailed eleven interacting residues which are conserved between the two bacteria and thus must be the most essential. Of these, most are basic amino acids (nine in *P. aeruginosa* and ten in *E. hormaechei*; Figure 6). Notable exceptions to this are Val62 and Leu93 in *P. aeruginosa,* and Tyr30 in *E. hormaechei*, as well as Asp75 in both pathogens. Some of the key residues placed at the α2 helix (positions 80, 83, 87, 89, 91, and 93) form an extended positive electrostatic potential surface, and this could play a role in driving the HPF into the ribosome or in the stabilization of its bound position. Even if they are not strictly functional homologs, the amino acids with identical locations in both bacteria are involved in interactions of the same nature. For example, in *P. aeruginosa,* His30 is involved in π-stacking and hydrophobic interactions with A784 and G1491, while in *E. hormaechei,* the identically located Tyr30 has these same interactions with A790 (Supp. Figures 4 A-B and 5 A-B).

**Figure 6.**
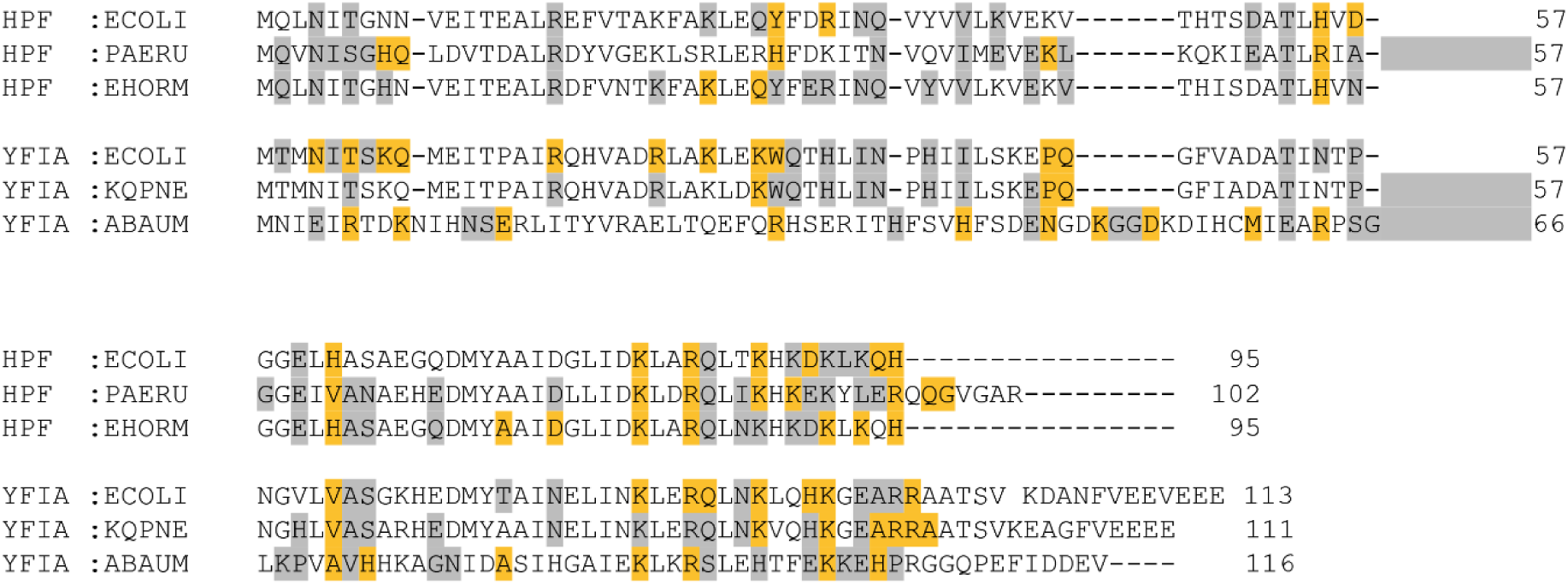
Illustration of the key amino acids from the hibernation factors HPF and YfiA that interact with the bacterial ribosomes in the present structures. The residues highlighted in orange are the most frequently in contact, while those in grey are less common. Sequence alignment was improved using pBlast with structural elements correspondence. *E. coli* structures used: PDB: 6y69 for HPF contacts with the *E. coli* ribosome; and PDB: 8fc1 for YfiA contacts with the *T. thermophilus* ribosome.

Many other amino acids also exhibit strong interactions with ribosomes in both bacteria (Figure 6). Interestingly, some residues specifically interact with modified nucleotides of 16S rRNA, a molecule which is crucial for ribosomal activity. One example is Thr53, which interacts with either m^2^G960 or m^2^G966 in *P. aeruginosa* and *E. hormaechei* (Supp. Figure 14). We also saw the conserved amino acid Arg83 contacting m^4^C1396 in *P. aeruginosa* and m^4^C1402 in *E. hormaechei* (Supp. Figure 12). Similarly, we observed *P. aeruginosa* Ile86 contacting m³U1492, while at an equivalent position in the *E. hormaechei* HPF, Asn86 contacts with m³U1498 (Supp. Figure 12).

Although both HPF proteins interact with the ribosome in very similar ways, they do also exhibit a degree of specificity. For instance, in *P. aeruginosa,* the residues Gln9, Gln97, and Gly98 are strong ribosome interacting partners but they are not present in *E. hormaechei,* and while residues Lys26 and Gln29 strongly interact with the *E. hormaechei* ribosome yet no equivalent interactors are found in *P. aeruginosa* (Figure 6).

### Overview of the HPF-ribosome interactions in Gram-negative bacteria

Most of the essential residues of HPF that interact with *P. aeruginosa* and *E. hormaechei* ribosomes have been already been described in *E. coli:* in dimers (100S ribosome); in the presence of the RMF protein (PDB 6h58, Beckert *et al*, 2018) as well as in monomers (70S ribosome) without RMF (PDB 6y69; Osterman *et al*, 2020). The position of HPF and its ribosome-binding site varies slightly across bacterial species, which results in differing lists of contacting residues. For instance, even with the high sequence identity between *E. hormaechei* and *E. coli*, the absence of an interaction between His8 and nucleotide A964 (next to modified m^2^G966) is highly notable in *E. coli,* where this histidine residue is replaced by asparagine (Figure 6). In the same way, His8 recreates this interaction together with Gln9 in *P. aeruginosa* (Supp. Figure 7A). Similarly, Lys26 and Gln29 residues of both *E. hormaechei* and *E. coli* interact with the phosphate groups of their neighboring residues m^3^U1498 (C1496 and G1497), whereas it is only His30 that performs this in the *P. aeruginosa* structure (Supp. Figures 5 A-B, and 12).

We also observed that the pair formed by His55 (Arg55 in *P. aeruginosa*) and Glu60 always interacts strongly with the C-terminal end of the protein uS9 carboxylate terminal. (Supp. Figure 9). In addition, in *P. aeruginosa,* Glu60 also in contact with nucleotide m^2^G960 (Supp. Figure 14). Finally, the strictly conserved basic residues of the second half of the α2 helix are exactly superimposable in three dimensions onto the Gram-negative bacteria, and they form the same interactions with the same nucleotides, regardless of the bacteria: base or base/phosphate contacts for Lys80 and Lys89; and base or sugar/base contacts for Arg83, Lys87, and Lys91. Moreover, residues located at positions 53 (T53) and 62 (V62/H62) in *P. aeruginosa* and *E. hormaechei* forms an invariant contact motif with m^2^G960/m^2^G966 (Supp. Figure 7 A-B). This motif seems highly specific to HPF, and can be further stabilized by other residues such as Ile40, Glu51, and Glu60 in *P. aeruginosa*. As illustrated here, these bacteria feature several highly conserved HPF residues which play an essential role in ribosomal binding.

### YfiA interactions with *K. quasipneumoniae* and *A. baumannii* ribosomes

The structure of the YfiA protein also exhibits a (β - α - β - β - β - α) fold identical to that of the HPF protein, but its C-terminal part has a tail which extends longer than that of HPF (Supp. Figure 1). As in previous studies, the C-terminal end of YfiA could not be modeled, as no clear density was observed in the map (residues 100-113 in *K. quasipneumoniae* and 105-116 in *A. baumannii*). Considering the high overall quality of the map, this suggests that the very end of the C-terminal tail is extremely flexible and probably does not tightly interact with the ribosome. The YfiA protein and its rigorously conserved fold occupies the same position as the HPF protein within the ribosome (Figure 2). It interacts with 16S rRNA and with uS7 and uS9 ribosomal proteins in both *K. quasipneumoniae* and *A. baumannii* (Table 1). The YfiA proteins of these two bacteria interact mainly with the C-terminal end of the uS9, with amino acids straddling the β3 and β4 strands, as observed in our HPF structures (see zone 7 in Figures 7 and 8, and Supp. Figure 9 C-D). For uS7, the contact is achieved by the C-terminal extremity of YfiA in both organisms (see zone 8 in Figures 7 and 8, and Supp. Figure 10 C-D).

**Figure 7.**
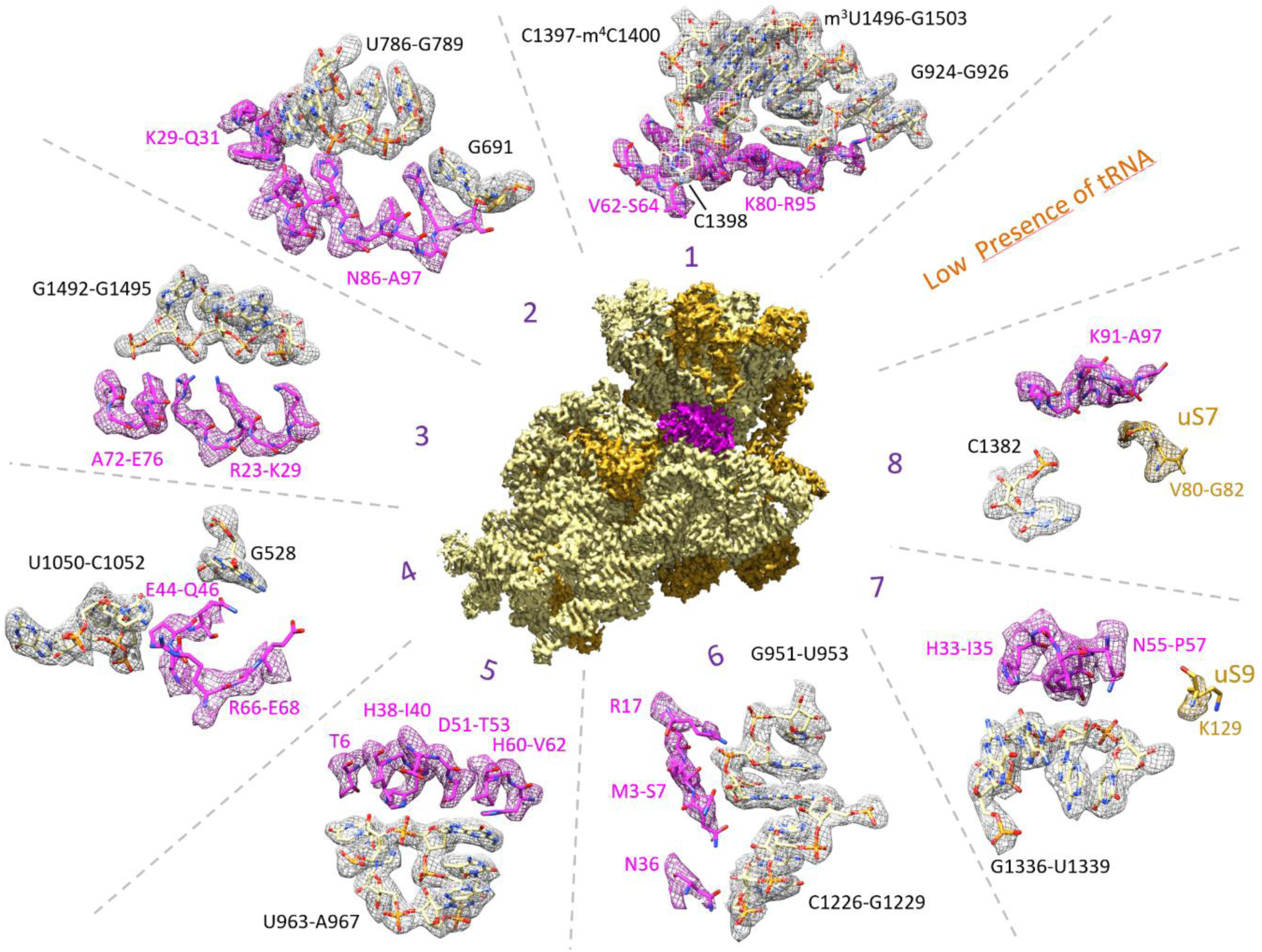
Interactions of the hibernation factor YfiA with the *K. quasipneumoniae* ribosome. To aid in visualization, the interaction spots are presented in zones numbered 1 through 8, with an extra area for tRNA. The nucleotide tracks of ribosomal 16S rRNA involved in binding to YfiA are black, the amino acids contacting YfiA are magenta, ribosomal proteins uS7 and uS9 are gold, and tRNA nucleotides are orange.

**Figure 8.**
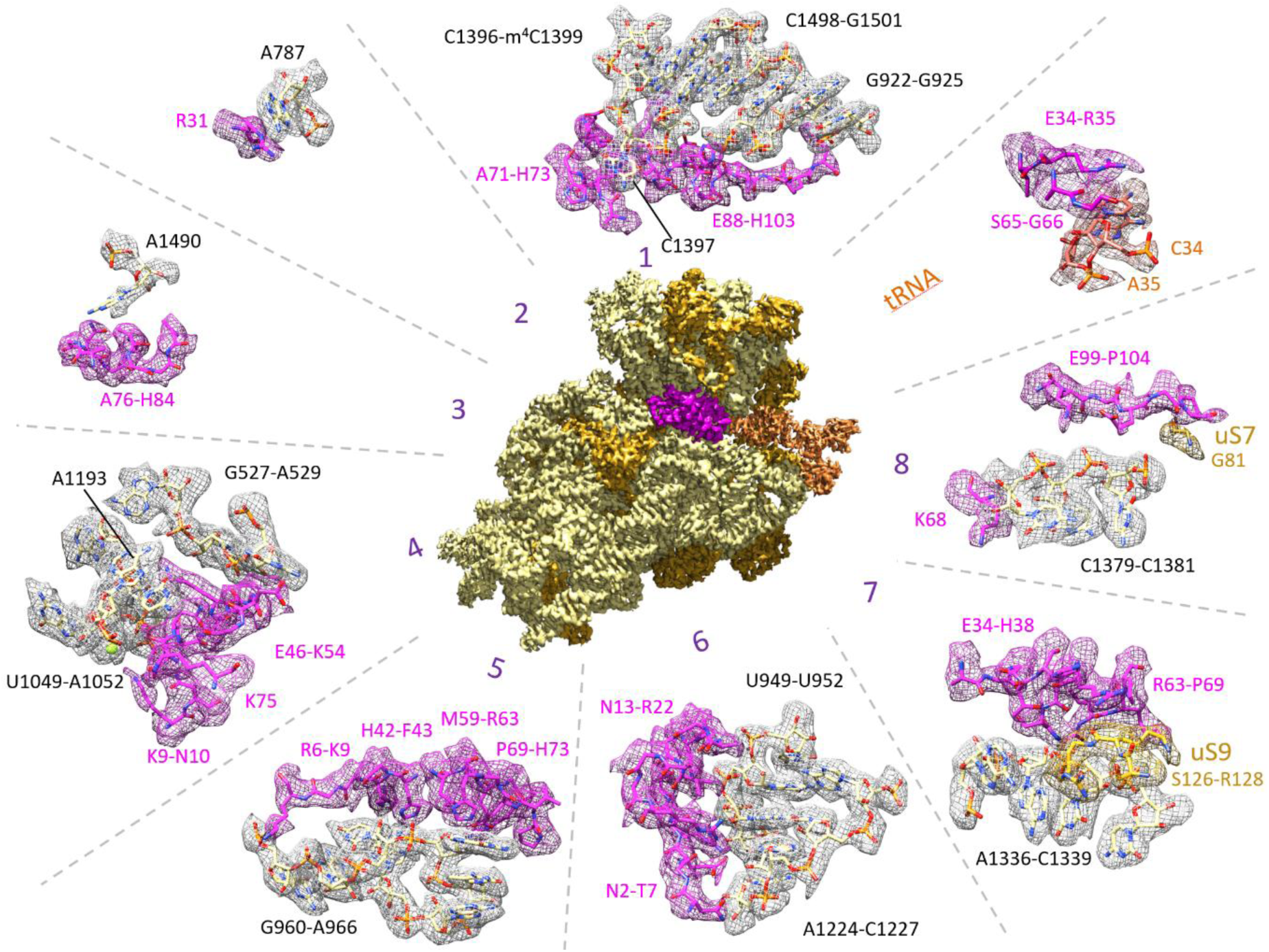
Interactions of the hibernation factor YfiA with the *A. baumannii* ribosome. To aid in visualization, the interactions are partitioned geometrically into zones 1 through 8 along with a highlighted tRNA zone (orange) that highlights the interaction between HPF and the E-site tRNA. The nucleotide tracks of ribosomal 16S rRNA involved in binding to YfiA are black, YfiA-contacting amino acids are magenta, ribosomal proteins uS7 and uS9 are gold, and tRNA nucleotides are orange.

In both bacteria, YfiA interacts with 16S rRNA in a similar manner, and much as HPF does. The ribosomal regions interacting with the YfiA mainly involve helices h28, h30, h31 and its stem loop, h44, and the h42/h29 junctions (Table 1). Despite the common fold, both bacteria do have their own specificity: the YfiA of *K. quasipneumoniae* strongly interacts with h23 and h24 and their stem loop, while that of A. *baumannii* strongly interacts instead with the h43/h28 junction (Table 1). In both bacterial YfiA proteins, an extended contact motif is observed at the center of the β-sheet (in the center of each strand β2, β3, and β4), and this motif is identical to that observed for HPF in our structures. As with HPF, YfiA strongly interacts with several modified 16S nucleotides. The three residues of YfiA from *K. quasipneumoniae* (Thr53, His60, and Val62) interact with the modified m^2^G964 (Supp. Figure 13 A), while the six *A. baumannii* YfiA residues interact with m^2^G963 (Supp. Figure 13 B). The arginines Arg83 (*K. quasipneumoniae)* and Arg92 (*A. baumannii)* interact with modified nucleotides m^4^C1400 and m^4^C1399 (Supp. Figure 13 A-B). Finally, we also observed an interaction between Asn86 and m^3^U1496 in *K. quasipneumoniae* (Figure 7 zone 1 and Supp. Figure 13 A), while no equivalent interaction was observed for *A. baumannii*.

The locations of the essential contacts between YfiA and its secondary structure elements are almost identical to those observed for HPF-bound ribosomes, with the exception of the long β2-β3 loop in *A. baumannii* YfiA which presents a rich interacting pattern involving Glu46, Asn47, and Lys50 to Asp53 (Supp. Figure 13 B). The YfiA hibernation factors found in these two pathogens exhibit significant differences in their primary sequences, despite having similar lengths (111 aa and 116 aa, respectively; Figure 6), which makes the proper alignment of these sequences using classical tools quite difficult. By combining multisequence and structural alignments, including the position and length of alpha helices and beta strands, we can see that the ribosomal interaction networks created by YfiA in these two bacteria are similar (Figure 6). Among the essential amino acids of the YfiA protein in *K. quasipneumoniae,* few are substituted by histidine in *A. baumannii*, resulting in a His-rich set of interacting residues similar to that observed with HPF. This may explain why there has been some confusion about the identity of *A. baumannii* YfiA, incorrectly called ‘HPF’ in 7m4z (Zhang *et al*, 2021). The track of basic amino acids located on one side of the α2 helix is very similar to the one observed in HPF, and involves the positions 80, 83, 87, 91, and 95 in *K. quasipneumoniae,* and 89, 92, 96, 100, and 103 in *A. baumannii.* The remaining non-basic residues which are strongly associated with the ribosome in *K. quasipneumoniae* include in: Trp30 in contact with A788 (Supp. Figure 4 C); Ile40 with U963 (Supp. Figure 7 C); Val62 with m^2^G964 and C1398; and Ser64 with C1398 (Supp. Figure 3 C). Meanwhile, in *A. baumannii*, the non-basic strongly associated residues are Ala71 which is in contact with m^2^G963 and C1397, and Asn78 contacting A1490 (Supp. Figure 13 B). In a similar fashion, the basic residues Arg31 (contacting A787), His42 (contacting U962, m^2^G963), and His73 (contacting m^2^G963, C1397) in *A. baumannii* play an equivalent role to that of the non-basic residues Trp30, Ile40, and Ser64 in *K. quasipneumoniae* (Supp. Figure 13 A-B). Furthermore, interaction patterns are sometimes reproduced in a manner that goes beyond simple residue-to-residue equivalence, as seen in the Lys29–Trp30/Arg31 group, where hydrophobic contributions play the dominant role (Figure 13). We also observed certain interactions specific to a bacterial species. In *K. quasipneumoniae* YfiA, the strong ribosome interactors are Arg95, Arg96, Ala97 with Asn75 contacting the 16s rRNA and Arg96 additionally interacting with ribosomal protein uS7 (Val80, Gly81, Gly82; Supp. Figure 10 C). In *A. baumannii,* its specific strong contacting amino acids include Glu15, Lys50, Asp53, and Ala81 (Figure 6). In addition, Met59 interacts with the methylated nucleotide m^2^G963, and Arg63 with the uS9 protein (Lys127, Arg128; Figure 6 and Supp. Figures 9D and 13B). It is also notable that in *A. baumannii* both His42 and His73 interact with the modified nucleotide m^2^G963 (Supp. Figure 13B), while no equivalent interaction was observed in *K. quasipneumoniae*.

We did also detect some weaker amino acid interactions which occur at equivalent positions in both *K. quasipneumoniae* and *A. baumannii*, but their roles and functional equivalences were not always clear. As we explain here, the significant differences found in the primary sequence of *A. baumannii* did not alter the ribosomal interaction motif.

### Overview of the interaction of YfiA with the ribosome in Gram-negative bacteria

The ribosome-binding mode of the YfiA protein has been studied in *E. coli*, but some of these studies were based on hybrid systems combining *E. coli* hibernation factors SHPF and RMF with *T. thermophilus* ribosomes (Chen *et al*, 2023; Polikanov *et al*, 2012). Nonetheless, the obvious similarity between the interaction networks observed in our *K. quasipneumoniae* YfiA-bound ribosomal structure and that of *E. coli* YfiA bound to *T. thermophilus* ribosomes can be further explained by their closely related primary sequences, as they share 91% identity. Thus, any comparison between *K. quasipneumoniae* YfiA and *E. coli* YfiA ribosomal interactions is relatively straightforward. However, the primary sequences of the hibernation factors in *A. baumannii* are significantly different than those of *E. coli* (32.63%) and *K. quasipneumoniae* (33.68%), and this may have led to a confusion in the naming conventions used in the literature, with one hibernation factor being labeled as the other.

We compared the YfiA-bound *A. baumannii* ribosomes in our study with the previously deposited structure (PDB 7m4z; (Zhang *et al*, 2021)) and found that the YfiA proteins were greatly similar, with just a few notable differences in positioning around the ribosomal small sub unit (SSU). Essential residues found in both structures include Arg6, Lys9, Glu15, Arg31, His42, Lys50, Arg63, Lys68, Pro69, Ala71, Val72, His73, Lys89, Arg92, His96, Lys100, and Lys101. That said, the list of ribosome residues interacting with YfiA are slightly different in our cryo-EM structure, indicating that there is a relative flexibility in the binding mode of the proteins promoting hibernation. For instance, Asn78 and His103 interact with the ribosome in our structure, but this was not observed in the previously published one. Similarly, we observed nucleotides A529, G925, U1049, C1379, m^4^C1399, A1490, A1500, and uS7 ribosomal protein (G81) interacting with YfiA, but they are not present in Zhang *et al* (Zhang *et al*, 2021). It is surprising that the interactions with modified nucleotides m^4^C1399 and m^2^G963 conserved in all of our structures were not mentioned in Zhang *et al*.(Zhang *et al*, 2021).

### Presence or absence of tRNA in the ribosomal E-site

There has been a long debate about whether YfiA, due to its long C-terminal extremity, actually prevents tRNA from binding in the ribosomal E-site. We constructed our *in-vitro* complexes by incubating the purified ribosomes with tRNA^fMet^ at a 1:2 ratio of 70S to tRNA^fMet^. Interestingly, we found tRNA in the E-site of three out of the four bacterial hibernating ribosomes examined (Supp. Table 1). When, YfiA was present, E-site tRNA was found in 80% of hibernating ribosomes from *A. baumannii*, but only 20% of those from *K. quasipneumoniae* (Supp. Table 1 and Supp. Figure 15). In the presence of HPF, E-site tRNA was found in 81% of hibernating ribosomes from *E. hormaechei*, but none at all were found in *P. aeruginosa* (Supp. Table 1 and Supp. Figure16). In fact, there is no correlation between the presence of E-site tRNA and the percentage of hibernating 70S ribosomes (Supp. Table 1). Furthermore, our results suggests that the presence or absence of tRNA depends on the bacterial species rather than the type of hibernation factor, although we do not have a clear explanation for why this might be.

Similarly, previous studies show that the C-terminal extremity of HPF interacts with E-site tRNA (Korostelev *et al*, 2006; Selmer *et al*, 2006; Yusupov *et al*, 2001). When tRNA was found in our hibernating ribosomes, it reacted with the ribosome in the same canonical ways as previously described for the 30S and 50S subunits (Supp. Figure 17).

### Specific observations near the mRNA channel, including the presence of bS21

In the structures we present here, we did not observe the hibernation factor RMF, principally because we analyzed the 70S ribosomal fraction. However, it is still intriguing that *E. hormaechei* ribosomes hibernates with the hibernation factor HPF, even when YfiA is still present. In our detailed analysis, we observed hibernating ribosome structures of varying compositions, and identified four distinct configurations (Supp. Figure 15) of YfiA-induced hibernating ribosomes from *K. quasipneumoniae* and *A. baumannii*. The first class contains hibernating ribosomes devoid of both tRNA and the bS21 protein, with mRNA forming a double helix with the anti-Shine-Dalgarno (SD) sequence at the 3′ end of the 16S rRNA. In the second, we identified structures where tRNA is present, and the double helix with the anti-SD sequence is also distinctly visible. In the third group, we observed hibernating ribosomes containing only bS21 and no E-site tRNA or mRNA, and therefore without the SD/anti-SD double helix. In the last class, hibernating ribosomes contain both E-site tRNA and bS21, but no mRNA (Supp. Figures 15 A-B). As for the ribosomes put into hibernation by HPF, we identified two conformational states each (Supp. Figure 16) for *P. aeruginosa* and *E. hormaechei*. In *P. aeruginosa,* one class consists of hibernating ribosomes containing bS21, while the other contains ribosomes lacking bS21 but containing mRNA, with the SD/anti-SD double helix clearly visible. Interestingly, we did not observe any E-site tRNA in these hibernating ribosomes (Supp. Figure 16 A 1-2). In contrast, no density corresponding to bS21 proteins was detected in our reconstructions in *E. hormaechei*. Both hibernating confirmations for this pathogen show a clearly defined mRNA–anti-SD helix at the 3′ end of the 16S rRNA, and while the second class features bound E-site tRNA, this is absent in the first.

## DISCUSSION

To identify the stalled 70S ribosomal complex in the presence of non-stop mRNA, tRNA^fMet^, and *trans*-translation factors tmRNA and SmpB (Guyomar *et al*, 2021), we performed cryo-EM and single particle analysis on 1.11 to 1.66 million particles collected from the 70S ribosomes of four ESKAPE pathogens (see Materials and Methods). Remarkably, we were able to isolate clusters of ribosomes bound with hibernation factors in all of them, at rates of approximately 8% to 26% (Supp. Table 1). Since they do not allow for tmRNA-SmpB binding, these hibernating ribosomes were probably formed prior to the addition of those *trans*-translation partners, due to the intermediate cold temperature step of our ribosome purification process (see Material and Methods).

For each of the bacteria studied, we unambiguously detected a unique hibernation factor in the ribosome: HPF in *P. aeruginosa* and *E. hormaechei,* and YfiA in *K. quasipneumoniae* and *A. baumannii*. With the exception of *P. aeruginosa* (with no known YfiA homolog, see Figure 3), the resolution of the cryo-electron density maps was sufficient to discriminate between the HPF and YfiA sequences, and thus to unambiguously identify the hibernation factors present in the bacteria (Supp. Figure 2). Interestingly, in these new structures there were no density traces to indicate the presence of RMF, which is the hibernating factor known to be crucial for forming 100S dimers in *gamma-*proteobacteria (Polikanov *et al*, 2012; Maki & Yoshida, 2022). In the disome of the *E. coli* structure, RMF is situated near the decoding center of the 30S subunit within a cavity formed by the ribosomal proteins uS2, uS7, uS9, and uS21(Maki & Yoshida, 2022; Beckert *et al*, 2018). This position allows RMF to interact with the regions surrounding the mRNA channel and the anti-Shine-Dalgarno (aSD) site (PDB 6h4n; (Beckert *et al*, 2018)). We superimposed that structure on our maps, and could thus confirm that the previously described binding sites for HPF and RMF also align with our structures (Supp. Figure 18). During this analysis, we made an additional observation about the relationship between the C-terminal region of the hibernating factor (either HPF or YfiA) and the RMF binding position. Although we were not able to resolve the last 12 residues of the YfiA C-terminal tail, we were still able to see that this tail progresses towards the binding site of RMF, and that this progression is not very clear in the tail of short HPF. This might explain why RMF is not visible in our YfiA-bound cryo-EM maps, but it is quite intriguing nonetheless to observe 70S ribosomes hibernating with HPF. Alternately, since our analysis focused solely on the 70S ribosome fraction without the 100S fraction, maybe RMF binding is less clear as previously thought. Or maybe 70S ribosomes can hibernate with HPF under certain conditions by following a different pathway when RMF is absent.

It would be interesting to explore why different hibernation factors are used to perform similar tasks, and why one is chosen over the other. The answers to these questions may lie in the adaptations of various bacteria to specific environmental challenges, as previously described for other organisms (Nicholson *et al*, 2022; Barandun *et al*, 2019). For example, the third class of bacterial hibernation factors, the Balon family, has unique binding properties in *P. urativorans* which suggests that a specialized mechanism was likely tailored to that organism’s lifestyle or environmental niche (Helena-Bueno *et al*, 2024). Unlike the more well-known hibernation factors RMF and HPF, Balon binds not just to inactive ribosomes but also to those actively engaged with mRNA and tRNA, and this indicates the existence of a unique form of translational regulation under stress conditions (Helena-Bueno *et al*, 2024).

### Hibernation factors HPF and YfiA interact with modified nucleotides

Of the 95 amino acids shared between the HPF and YfiA hibernation factors in *E. coli*, 35 are rigorously identical. Only four of these, the basic residues Lys80, Arg83, Lys87, and Lys91, are involved in ribosomal interactions, thus the others must contribute to conserving the protein fold. In comparing the ribosome-contacting hibernation factor residues in our four bacterial models, we were able to identify certain residues at conserved positions as being essential (Figure 6). Although the β4 strand and α2 helix are commonly involved in both HPF and YfiA ribosome binding, in HPF the β1 strand also contributes additional ribosomal contacts. Similarly, the sequence is weakly conserved between positions 60 and 64 (using *E. coli* numbering as a reference), but this does not seem to affect the conservation of an interaction motif.

Depending on the bacterial species and their respective hibernation factors, the number of essential interacting residues varies, yet the equivalence we establish here highlights the strong interactions with the modified nucleotides m^2^G966, m^3^U1498, and m^4^C1402 (again using *E. coli* numbering). Specific interactions of YfiA with these modified nucleotides of 16S rRNA have already been described for *E. coli* YfiA in the *T. thermophilus* ribosome, where Thr53, Val60, and Val62 are known to interact with m^2^G966, Arg83 interact with m^4^C1402, and Asn86 with m^3^U1498 (Polikanov *et al*, 2015). In both of our YfiA and HPF structures, we observed a strong interaction platform for the recognition and binding of the modified guanine m^2^G966 (*E. coli* numbering). The amino acids involved form a highly conserved patch on the β3 and β4 strands (Supp. Figure 12): residues Glu51, Thr53, and Val62 for *P. aeruginosa* HPF; Thr53, and His62 for the *E. hormaechei* HPF; Thr53, His60, and Val62 for the *K. quasipneumoniae* YfiA; and Glu61, Pro69, and Ala71 for the *A. baumannii* YfiA. These amino acids are sometimes supported by additional residues, including His42, Met59, and His73 in *A. baumannii,* and Ile40 in *P. aeruginosa*. The interaction network organized around the nucleotide m^4^C1402 (*E. coli* numbering) is also highly conserved, and it involves a single essential arginine: Arg83 in *P. aeruginosa, E. hormaechei* and *K. quasipneumoniae;* and Arg92 in *A. baumannii*. As for the third nucleotide, m^3^U1498 interactions are maintained with Ile86 in *P. aeruginosa* and with Asn86 in *E. hormaechei* and *K. quasipneumoniae,* but no such interaction is observed in *A. baumannii* (Supp. Figures 12 and 13).

The interactions between HPF/YfiA and these three highly conserved modified nucleotides appears to be an inherent property of the RaiA family of hibernation factors. The interactions allow these hibernation factors to achieve several important things simultaneously; they prevent translation initiation by blocking the m^2^G966-m^4^C1402 structural “clamp” which would normally stabilize P-site tRNA; they specifically protect correctly folded ribosomes against RNase degradation by recognizing two essential late modifications m^4^C1402 and m^3^U1498 serving as checkpoints for 30S maturation; and they prevent the binding of certain 30S-targeting antibiotics such as the aminoglycosides streptomycin, paromomycin, and gentamycin.

### Hibernation states as a drug target

Understanding the mechanisms of resistance and pathogenicity is crucial for developing strategies to combat infectious diseases. When combined with previously published cryo-EM studies, this study offers a comprehensive molecular-level description of how hibernation factors induce ribosome dormancy by binding on active sites and interacting with modified nucleotides (Polikanov *et al*, 2012; Syroegin *et al*, 2022). Furthermore, genetic and biochemical evidence from *E. coli* (Niven, 2004; Yoshida *et al*, 2002) and functional studies of *B. subtilis* (Akanuma *et al*, 2016) have demonstrated that ribosomal dimerization is crucial for bacterial survival during nutritional stress (the stationary phase) and for efficient regrowth after dormancy, thus emphasizing the physiological relevance of hibernation factors (Prossliner *et al*, 2018; Trösch & Willmund, 2019; Nissley *et al*, 2025; Prossliner *et al*, 2021).

Recent investigations of ribosomal structures bound to antibiotics have shown how these drugs inhibit translation by trapping the ribosomes in specific conformational states (Ekemezie & Melnikov, 2024; Syroegin *et al*, 2022). They have also revealed how resistance can emerge through mutation or modification of drug-binding sites (Canu *et al*, 2002; Gomez *et al*, 2017; Demirci *et al*, 2014; De Stasio *et al*, 1989; Hoffman, 2001). Interestingly, both HPF and YfiA overlap with the binding sites of various ribosome-targeting antibiotics, including neomycin, gentamicin, hygromycin B, capreomycin, tetracycline, tigecycline, and kasugamycin (Ekemezie & Melnikov, 2024; Khusainov *et al*, 2017).

The recently identified Balon protein, a novel ribosome hibernation factor, differs from previously characterized factors in that it binds to the ribosomal A-site and contacts both the decoding center of the small subunit and the peptidyl transferase center of the large subunit (Helena-Bueno *et al*, 2024). Similar to HPF and YfiA proteins, Balon’s binding position also overlaps with the sites typically occupied by several antibiotics, including thermorubin, hygromycin B, amikacin, neomycin, and gentamicin. A similar principle was observed in *B. subtilis*, where BsHPF (long form of HPF) engages overlapping ribosomal sites and prevents antibiotic binding, emphasizing the conserved nature of this mechanism (Beckert *et al*, 2017) These findings highlight the complex dynamics between hibernation factors and antibiotic binding within the ribosome, the exploration of which has provided insights into potential mechanisms of antibiotic resistance as well as ribosomal protection.

Currently, inhibiting translation is one of the most common antibiotic modes of action for targeting active ribosomes and its active sites (Lin *et al*, 2018; Parajuli *et al*, 2021). Hibernation factors seem to play a vital role in protecting the ribosome from these molecules and from RNase, allowing bacteria to wait for the storm to pass (Prossliner *et al*, 2018; Ekemezie & Melnikov, 2024). As such, the hibernating state of ribosomes under specific stressful conditions would be an interesting target for new antimicrobial compounds. Our high-resolution structures of bacterial ribosome complexes and the accompanying hibernation factors have mapped precise binding pockets and interaction surfaces, thereby revealing possible drug-target sites as well as binding motifs that could be targeted by small molecules or peptides. For example, inhibitors designed to block hibernation factor binding or inhibit their contacts at the decoding center would be able to prevent ribosome hibernation, forcing bacteria to continue active translation or get rid of the bound proteins, thus increasing their susceptibility to antibiotics.

## CONCLUDING REMARKS

Since protein synthesis is the most energy-consuming process in cells, the prevention of ribosomal degradation could be evolutionary favorable, as it allows cells to maintain steady survival based on the available resources. This is particularly beneficial when nutrients are limited, and enables rapid resumption of growth when they become available again. This ability to quickly resume growth is crucial for survival and competitive fitness. Given that at any moment approximately 60% of the microbial biomass is in a dormant state, ribosome hibernation factors are probably the most prevalent ribosomal partners within bacterial cells. Although most of these dormant ribosomes are bound by hibernation factors, most research on how antibiotics work has concentrated on actively translating ribosomes that lack such factors. A more detailed understanding of ribosome hibernation will allow us to further understand cellular stress responses and the regulation of protein synthesis. In turn, this understanding will lead to potential applications in medicine and biotechnology, such as the development of strategies to combat bacterial persistence and improve the efficacy of antibiotics against highly pathogenic bacteria. According to a World Health Organization report in 2021, nine out of fifteen species in the bacterial priority pathogens list are Gram-negative(World Health Organization. Bacterial priority pathogens list to guide research). Given the anticipated rise in antimicrobial resistance, there is an urgent need to explore new therapeutic avenues such as dormant-ribosome targeting if we are to successfully combat this growing challenge. This structural study of hibernating ribosomes in four highly pathogenic Gram-negative bacteria provides meaningful initial insights, and we hope that this work paves the way for structure-based strategies which will combine computational modeling, virtual screening, and experimental biochemical testing in order to design new compounds that will be able to interfere with ribosome hibernation.

## MATERIALS AND METHODS

### Strain growth conditions

Wild-type (WT) strains of *Klebsiella quasipneumoniae (K. pneumoniae,* DSM 26371 / ATCC 700603)*, Acinetobacter baumannii* (ATCC BAA1605), and *Pseudomonas aeruginosa PA14* (DSM 19882) were grown at 37 °C in tryptic soy broth (TSB) media, while *Enterobacter hormaechei subsp. Xiangfangensis (Enterobacter cloacae* complex, ATCC BAA2341) was grown in brain heart infusion (BHI), and *Escherichia coli* in Luria-Bertani (LB) medium. Where necessary, the media were supplemented with 1.5% agar (w/v) and/or antibiotics. Unless stated otherwise, chemicals and oligonucleotide primers were purchased from IDT / Sigma-Aldrich.

### Ribosome isolation and purification

Ribosomes were purified from WT strains of *K. quasipneumoniae*, *A. baumannii, P. aeruginosa PA14,* and *E. hormaechei*. From each fresh plate of bacteria, a colony was inoculated into 50 mL TSB or BHI and grown overnight (37 °C and 150 rpm). The following day, the cultures were diluted in 2 L of TSB or BHI medium to reach an initial absorbance of approximately 0.05 (OD_600_), then grown again (37 °C and 150 rpm). Cells were then harvested by centrifugation (4 °C at 10,000 rpm for 15 – 20 min.) at an absorbance of 0.3-0.5 (OD_600_). Pellets were washed in LB and centrifuged again at (4 °C at 14,000 rpm for 10 min), after which the bacterial pellets were stored at -80 °C until needed for purification of the ribosomes.

On the day of ribosome purification, bacterial pellets were thawed on ice for 30-40 min. The thawed pellets were resuspended for 15 min. at 30 °C in shaking conditions in a lysis buffer containing phenylmethylsulfonyl fluoride (1 mM PMSF, 20 mM Tris-HCl pH 7.5, 50 mM MgOAc, 100 mM NH_4_Cl, 1 mM EDTA, and 1 mM DTT, with the addition of 0.25 mg/mL lysosymes for the *A. baumannii* and *P. aeruginosa* strains). The pellets were lysed using a FastPrep (speed 6 for 30 sec., repeated 3 times), and treated with DNase I by incubating on ice for 10 min, then centrifuged (4 °C at 14,000 rpm for 15 min.) to remove any cellular debris. The supernatant was filtered through a sterile filter membrane (0.22 μm). At the same time, a sucrose cushion buffer was prepared (20 mM Tris-HCl pH 7.5, 50mM MgOAc, 100 mM NH_4_Cl, 0.5 mM EDTA, 1 mM DTT, and 1.1 M sucrose). The supernatant containing ribosomes was transferred with an equal amount (1:1 volume ratio) of sucrose cushion buffer into an ultracentrifugation tube. Ribosomes were isolated by ultracentrifugation (4 °C at 30,000 rpm for 20 h), then the supernatant was gently removed by inverting the tube and washing the pellets carefully twice with “TC Buffer” (10 mM Tris-HCl pH 7.5, 4 mM MgOAc, 50 mM NH_4_Cl, 0.5 mM EDTA, and 1 mM DTT). The ribosomal pellets were then dissolved in TC Buffer on ice for 6-7 hours, after which the pellets were loaded onto buffered sucrose cushions (10 mM Tris-HCl pH 7.5, 10mM MgCl_2_, 50 mM NH_4_Cl, 0.5 mM EDTA, 1 mM DTT, and 10%–50% [w/v] sucrose) to resolve 70S ribosomes by ultracentrifugation (4 °C at 23,000 rpm for 18 h) in an SW28 rotor (Beckman Coulter). The ultracentrifuged sucrose gradients were further fractionized into 500 µL aliquots using an ÄKTAprime fraction collector, which was also used to measure the absorbance of (OD_260_) fractions. The fractions containing 70S ribosome peak were pooled and further concentrated in a final centrifugal step (4 °C at 30,000 rpm for 20 h) with ribosome buffer (10 mM Tris-HCl pH 7.5, 10mM MgCl_2_, 50 mM NH_4_Cl, 0.5 mM EDTA, and 1 mM DTT). On the last day of the process, the supernatant was gently removed by inverting the tube. The ribosomal pellets were allowed to dissolve on ice in a minimal amount of ribosome buffer for overnight. Once they were completely dissolved, the ribosomal concentrations were measured and the ribosomes were immediately flash-frozen in liquid nitrogen and conserved at -80 °C for future use.

### *In vitro* accommodation complex and cryo-EM grid preparation

In order to obtain the mostly accommodated *trans*-translation state ribosomes possible, we prepared only the accommodated state complex, as previously described (Guyomar *et al*, 2021). This contains only the major partners of *trans*-translation (ribosomes, tRNA^fMet^, a short non-stop mRNA, tmRNA, and SmpB), purified from WT bacteria, and in *in vitro* conditions as close as possible to native conditions. Once the complexes were prepared, the concentration of 70S was adjusted to 140 nM in “Buffer III” (5 mM HEPES-KOH pH 7.5, 10 mM MgOAc, 50 mM KCl, 10 mM NH_4_Cl, and 1 mM DTT), then samples were directly applied to glow-discharged holey c-flat 4/2 and 2/1 µm grids. These grids were flash-frozen in liquid ethane using an EM-GP freezer (Thermo Fisher Scientific).

### Cryo-EM data collection

The flash-frozen grids were first investigated using our in-house TECNAI cryo-EM to check for tmRNA in order to confirm the presence of ribosomes in the *trans*-translation state. The grids were then transferred to a Titan Krios electron microscope (FEI) operating at 300 kV and equipped with an FEG electron source. Images were automatically recorded using EPU (Thermo Fisher Scientific) under low-dose conditions on a Gatan K3 direct electron detector, with a defocus range of -0.5 to -2.5 µm, a nominal magnification of ×81,000 and a final pixel size of approximately 1.1 Å. At the end of the acquisition, the datasets resulted in a total of 17,208 micrographs for *K. quasipneumoniae*, 15,401 micrographs for *A. baumannii,* 14,629 micrographs for *P. aeruginosa,* and 22,691 micrographs for *E. hormaechei* (Supp. Table 2).

### Cryo-EM data processing

CryoSparc (v4.3.1) (Punjani *et al*, 2017) was used to analyze the data. Briefly, the movies were dose-weighted and corrected for the effects of drift and beam-induced motion using Patch Motion Correction. The resulting aligned micrographs were also Fourier-cropped by a factor of 0.5, resulting in a final pixel size of 1.06 Å/pixel. Contrast transfer function (CTF) parameters were estimated using CryoSparc Patch-CTF. Electron micrographs showing signs of drift or astigmatism were manually discarded. This resulted in the retention of 16,996 micrographs for *K. quasipneumoniae,* 14,816 micrographs for *A. baumannii,* 14,493 micrographs for *P. aeruginosa,* and 15,273 micrographs for *E. hormaechei*. Particles were semi-automatically selected and extracted. To allow for efficient processing they were Fourier-cropped a second time by a factor of 0.5, resulting in images with 2.12 Å/pixel. The particles were then subjected to three rounds of 2D-classification in order to discard defective particles. This resulted in 1,411,532 good particles for *K. quasipneumoniae,* 1,565,495 for *A. baumannii*, 1,661,710 for *P. aeruginosa,* and 1,103,392 for *E. hormaechei*. After *ab initio* reconstruction and homo-refinement, the discrete heterogeneity present in each dataset was assessed using 3D classification without alignment. The separation of the datasets into ten discrete classes allowed us to identify some with hibernation factor containing ribosomes. After re-extracting the corresponding particles in boxes of 512 x 512 and returning to the original (non-super resolution) pixel size of 1.06Å/pixel, a second round of homogeneous refinement was done and followed by 3D classification into 5 discrete classes after subtracting the signal of the empty ribosome, and these results further improved the selection of hibernating ribosomes. Non-uniform refinement with optimized per-particle defocus and per-exposure-group higher-order aberration correction of these particles yielded maps for *K. quasipneumoniae* (at 2.3 Å)*, A. baumannii* (2.5 Å), *P. aeruginosa* (2.6 Å), and *E. hormaechei* (2.8 Å).

A final round of 3D classification was performed after subtracting the signal of the empty ribosome, and focusing in on a 40 Å-diameter sphere centered on either the E-site tRNA or on the RBS domain. This revealed populations of particles with or without E-site tRNA, and with either bS21 or the short mRNA. For *A. baumannii* this resulted in four groups: 106,853 particles with tRNA but without bS21; 74,087 particles with both; 26,517 particles with neither; and 17,801 particles with bS21 but lacking tRNA. Non-uniform refinement of the resulting populations yielded 4 separate maps at 2.7 Å, 2.7 Å, 2.9 Å, and 2.9 Å, respectively. For *P. aeruginosa* this resulted in only two populations: 50,773 particles containing bS21 and 81,011 particles without it, with no traces of E-site tRNA in either group. Non-uniform refinement of those two populations yielded two maps at 2.8 Å and 2.7 Å, respectively. For *E. hormaechei,* the non-uniform refinement also resulted in two populations, this time with 70,464 particles containing E-site tRNA and 16,968 particles without, and no trace of bS21 in either. Non-uniform refinement of those populations produced two maps at 2.8 Å and 3.1 Å, respectively. Finally, for *K. quasipneumoniae,* the refinement resulted in four separate populations: 251,038 particles containing neither tRNA nor bS21; 56,981 particles with tRNA but no bS21; 48,868 particles with bS21 but no tRNA; and 17,191 particles with both. Non-uniform refinement of those populations yielded maps at 2.5 Å, 2.7 Å, 2.7 Å, and 2.9 Å, respectively. Local resolutions were estimated using CryoSparc, and map quality was analyzed using Phenix mtriage software (Adams *et al*, 2010). The resulting density maps were then used for model building and refinement.

### Model building and refinement

UCSF-Chimera (Pettersen *et al*, 2004)was used to rigid-body fit the atomic model of the translating *E. coli* ribosome at 1.55 Å (PDB 8B0X) into the *K. quasipneumoniae* and *E. hormaechei* electron density maps, with each protein and RNA treated separately. The same approach was employed for *P. aeruginosa* using the previously published 2.6 Å cryo-EM structure (PDB 7unw), after removing the initiation facto IF2. While several structures already exist for *A. baumannii* (PDB 6V39 and 7M4Z), we decided to also start our modeling from the *E. coli* 8B0X structure, as close inspection of the published cryo-EM maps revealed a substantial error in pixel-size estimation. All models were then mutated according to their annotated genomic sequences and manually adjusted in the cryo-EM maps using COOT (v0.9.8.95) (Emsley *et al*, 2010). With the exception of pseudo uridines (which cannot be differentiated from uridines at our resolution), post-transcriptional modifications were only applied if visible in the electron density map and if the corresponding enzyme was annotated in the genome, and they were added based only on the genomic annotation. When needed, an atomic model of fmet-tRNA (taken for 8FTO) was rigid-body fitted into the electron density map, while the mRNA model was directly built onto the map using COOT. After the first real-space refinement of the whole ribosome, outliers were manually corrected in COOT. For each organism, Alfafold2 models of both HPF and YfiA were rigid-body fitted and real-space refined using COOT. Careful inspection of each hibernation factor residue to assess its fit into the density allowed for the unambiguous differentiation between HPF and YfiA. Using 8BOX as a reference, zinc, magnesium, and potassium ions were manually added into the remaining unmodelled blobs, but because of the final resolutions of the maps, we choose not to add water to the final model. Finally, the complete structures underwent five cycles of refinement in Phenix (v1.21.2-5419). Models were evaluated with MolProbity (Williams *et al*, 2018), and the remaining analyses and illustrations were done using UCSF-Chimera and ChimeraX (Goddard *et al*, 2018).

### Phylogenetic analysis

Complete protein sequences of ribosomal hibernation-promoting factors from *hpf* and *yfiA* genes in ESKAPE pathogens (*K. quasipneumoniae, A. baumannii, P. aeruginosa* and *Enterobacter spp.,* and *E. coli*) were retrieved from the NCBI GenBank database. Multiple sequence alignment was performed using MUSCLE v5.1 using the iterative refinement algorithm (UPGMB) to enhance alignment accuracy and conserve structural features (Edgar, 2004). Phylogenetic trees were constructed using IQ-TREE v2.2 and its maximum likelihood method, and the optimal substitution model was determined using ModelFinder (Kalyaanamoorthy *et al*, 2017). Branch support was evaluated through 1,000 ultrafast bootstrap replicates (UFBoot2) to ensure statistical reliability (Minh *et al*, 2020).

### Data availability statement

The structural models and maps generated in this study have been deposited to the Protein Data Bank with the accession numbers PDB ID-XXXX, PDB ID-XXXX, PDB ID-XXXX …. and to the Electron Microscopy Data Bank (EMDB) with the accession numbers EMD-XXXXX, EMD-XXXXX, EMD-XXXXX…..

## Acknowledgements

This work was supported by the Fondation pour la Recherche Médicale (Equipe FRM EQU202203014663) and the Agence Nationale de la Recherche (ANR-23-CE18-0031-01). The authors gratefully acknowledge the TEM^2^C (Transmission Electron Microscopy, Cryo&CLEM) facilities of Biosit UAR 3480 US18 (Univ Rennes, CNRS, Inserm) for the acquisition of preliminary cryo-EM data. The high resolution cryo-EM data were acquired at the NeCEN facility (Leiden University, the Netherlands), and funded by iNEXT-Discovery (Project number 2309-361 iNEXT - PID 27112). We are grateful to Ludovic Renault and the platform engineers for conducting the experiments and for their local support, and to Juliana Berland for her valuable feedback on the manuscript.

